# In silico analysis for potential proteins and microRNAs in Glioblastoma and Parkinsonism

**DOI:** 10.1101/2021.10.06.463376

**Authors:** Sayak Banerjee, Souvik Chakraborty, Tarasankar Maiti, Sristi Bisawas

**Affiliations:** Department of Physiology, Bhairab Ganguly College; Department of Zoology, Suri Vidyasagar College

**Keywords:** DifferentiallyF Expressed Genes, Glioblastoma, Parkinson’s Disease

## Abstract

In today’s world, neurodegenerative diseases such as Alzheimer’s disease, Parkinson’s Disease, Huntington’s Disease as well as brain cancers such as astrocytomas, ependymomas, glioblastomas have become a great threat to us. In this study, we are trying to find a probable molecular connection associated with two very much different diseases, Glioblastoma, also known as Glioblastoma Multiforme (cancers of microglial cells of our brain) and Parkinson’s disease. We at first downloaded the microarray datasets of these two diseases from Gene Expression Omnibus (GEO) and then analyzed them by the GEO2R tool. After analysis, we found 249 common upregulated differential expressed genes and 135 common downregulated differential expressed genes of these two diseases. Therefore the common differentially expressed genes, both upregulated and downregulated, were imported into STRING online tool to find out the protein-protein interactions. Now, this whole network was subjected to Cytoscape and the top ten hub genes were found by Cyto-Hubba plug-in. The top then hub genes are **EGFR, CCNB1, CDK1, CCNA2, CHEK1, RAD51, MAD2L1, KIF20A, BUB1**, and **CCNB2**. These all genes are upregulated in both diseases. To find out the biological processes, molecular functions, cellular components, and pathways associated with these hub genes Enrichr online software was used. We used miRNet software to determine the interactions of hub genes with microRNAs. This study will be useful in the future for drug targets discovery for these diseases.

## 1. Introduction

We’re looking for a possible biological link between Glioblastoma (brain malignancies affecting microglial cells) and Parkinson’s disease in this study. Glioblastoma, the most aggressive type of glioma, has extremely low survival rates, and diagnosis is based solely on brain imaging and biopsy sample analysis (Müller Bark et al., 2020). In a study, it was discovered that 0.73 to 4.49 people per 1,00,000 were diagnosed with glioblastoma (Grech et al., 2020). Glioblastoma can occur at any age, however, it is most common in adults between the ages of 55 and 60 (Hanif et al., 2017). Seizures, cognitive impairments, drowsiness, headache, fatigue, motor deficits, and other symptoms are common in people with glioblastoma, however, seizures and cognitive deficits seem to be the most common (IJzerman-Korevaar et al., 2018).

Parkinson’s disease (PD) is a neurodegenerative disease, affecting both upper motor and lower motor neurons, and ultimately leads to the impairment in motor movements. In PD, the dopamine level in our brain drops, and dopaminergic cells die, that’s why some PD patients experience cognitive impairments (Emamzadeh & Surguchov, 2018). The substantia nigra of the basal ganglia is primarily affected in Parkinson’s disease, and anatomical changes inside the cerebellum, such as contraction of the left cerebellum and reduction of grey matter in the right quadrangular lobe, are also observed (Prakash et al., 2016). According to one study, 1 to 2 people out of every 1000 can be impacted by Parkinson’s disease at any time (Tysnes & Storstein, 2017). Motor symptoms such as Tremors, bradykinesia, rigidity, postural instability, etc are common in Parkinson’s disease (Váradi, 2020).

In this study, we analyzed the datasets of Glioblastoma (GSE31262) and Parkinson’s Disease (GSE19587), obtaining from Gene Expression Omnibus, and found out hub genes and their pathways. We also studied the microRNAs associated with the hub genes.

## 2. Material and Methods

### 2.1 Data Selection

To obtain the datasets of these diseases we used the National Centre of Biotechnology Information (NCBI) online search tool Gene Expression Omnibus (GEO) (available in https://www.ncbi.nlm.nih.gov/geo/) (Clough & Barrett, 2016). We can find out the datasets by using disease name, GEO accession number, or even the author-name as search input. Here we used the disease name as search input. Inside the GEO we found several options like “Top Organisms”, “Study type”, “Attribute name” etc. In this study, we selected “Homo sapiens” inside the “Top Organisms” option, ‘tissues’ inside the “Attribute name” option, and ‘expression profiling by array’ inside the “Study type” option. Therefore we chose the dataset GSE31262 for glioblastoma and GSE19587 for Parkinson’s Disease (PD). We selected 9 glioblastoma patient samples and 5 control samples in dataset GSE31262, whereas we chose 12 PD patients and 10 control samples in dataset GSE19587 for analysis.

### 2.2 Differential Gene Expression and selection of Differentially Expressed Genes (DEGs)

We used the GEO2R tool (available at https://www.ncbi.nlm.nih.gov/geo/geo2r/) to analyze the two datasets GSE31262 for glioblastoma and GSE19587 for Parkinson’s Disease (Barrett et al., 2012). GEO2R enables us to compare two or more groups of samples (in this case patients and controls) in the GEO series to find out the differentially expressed genes. A text file with a list of differentially expressed genes was obtained after running the GEO2R tool. It contains gene IDs, adjusted p values, p values, t values, b values, LogFC values, gene symbols, and gene titles, in turn. To calculate the p value, the t test was used. No outliers were present in these datasets.

After downloading the lists of DEGs for these two diseases individually, they were subjected to different excel sheets. DEGs with a p value greater than 0.05 were deleted. In this study, we took the mod values of LogFC and its range is |0.5| to |7|. It indicates that DEGs with a positive LogFC value, are upregulated and with a negative LogFC value are downregulated in these diseases.

### 2.3 Identification of common DEGs

To find out the common DEGs in these two diseases, we used FunRich (version 3.1.3), a desktop-based tool (Fonseka et al., 2021). FunRich creates a Venn diagram through which we can easily identify the common DEGs. In this study, we created two Venn diagrams, one for common upregulated DEGs and another one for common downregulated DEGs in these two diseases.

### 2.4 Network Analysis

The list of common upregulated and common downregulated DEGs, both were imported to STRING (available in https://string-db.org/) to obtain the protein-protein network of DEGs. The protein-protein network, which we obtained in STRING, directly was subjected to Cytoscape from String (Shannon, 2003). Inside the CytoHubba, we used the cytohubba plugin to find out the top 10 hub genes (genes with maximum connectivity with other genes). Therefore **Enrichr** (available in https://maayanlab.cloud/Enrichr/) online software was used to enrich the **GO ontology (GO)** and **KEGG** pathway enrichment (Reimand et al., 2019). To find out the interactions between the hub genes and microRNAs, we imported the top 10 hub genes into the **miRNet** (available in https://www.mirnet.ca/) online software (Fan et al., 2016)

## 3. Results

In this study, we used the microarray datasets from Gene Expression Omnibus to identify the differentially expressed genes for glioblastoma (GSE31262) and Parkinson’s disease (GSE19587). Then we analyzed the DEGs, protein-protein network, found the hub genes and their interactions with microRNAs. We used GEO2R to identify the DEGs and to normalize the data.

In the dataset GSE31262, we found 2787 DEGs (1229 DEGs are upregulated and 1488 DEGs are downregulated) out of 32878 genes, and in the dataset GSE19587, out of 22280 genes, 3023 genes were differentially expressed (2051 DEGs are upregulated and 972 DEGs are downregulated), which passed out the set cut off of p-value 0.05 and mod value of LogFC (|0.5| to |7|).

The common differentially expressed genes were found by the Venn diagram using FunRich. Here, we found out common upregulated DEGs and common downregulated DEGs for these diseases. By using FunRich, we identified 249 common upregulated DEGs and 135 common downregulated DEGs in these diseases (Figure-2)

**Fig. -1:**
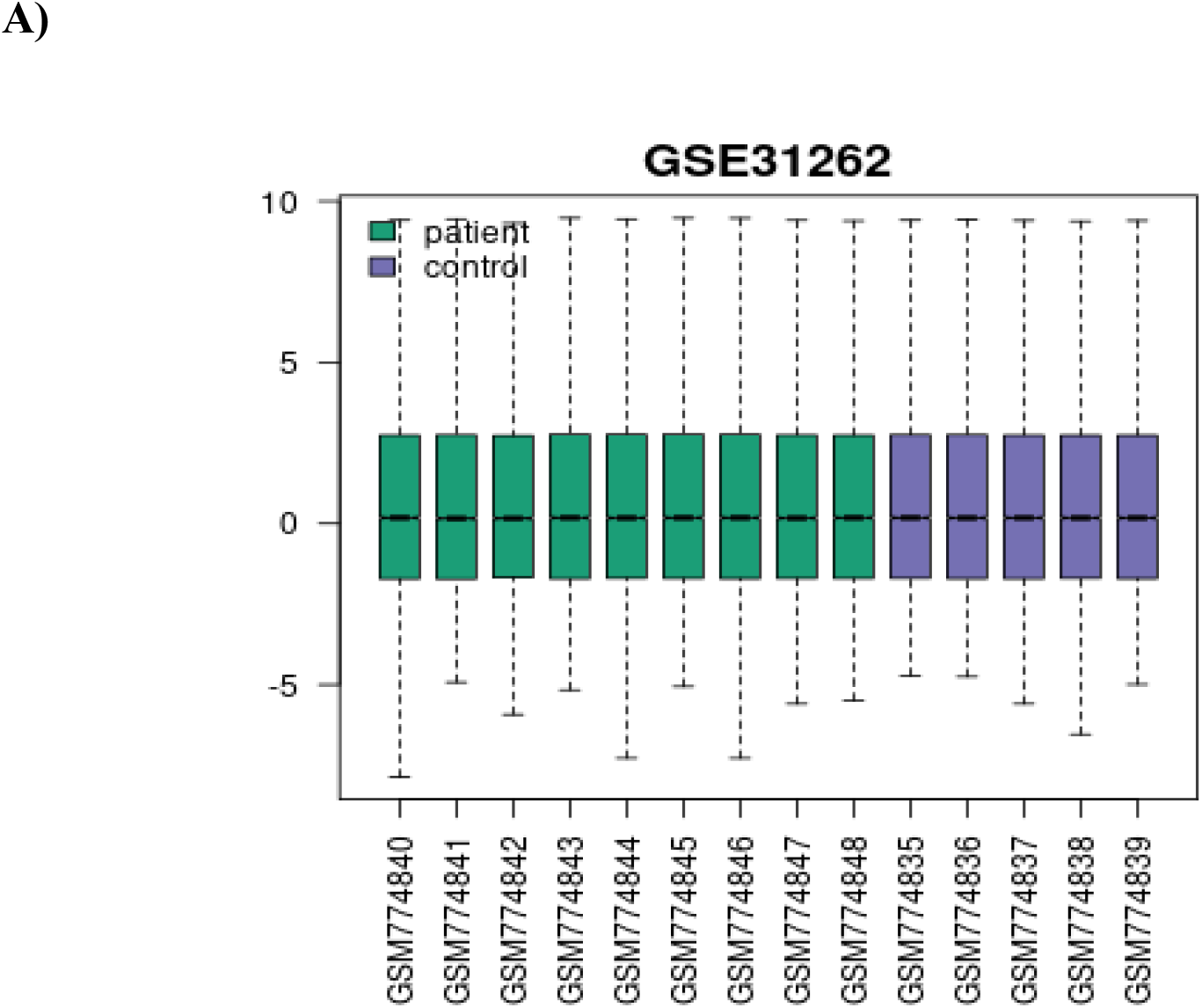
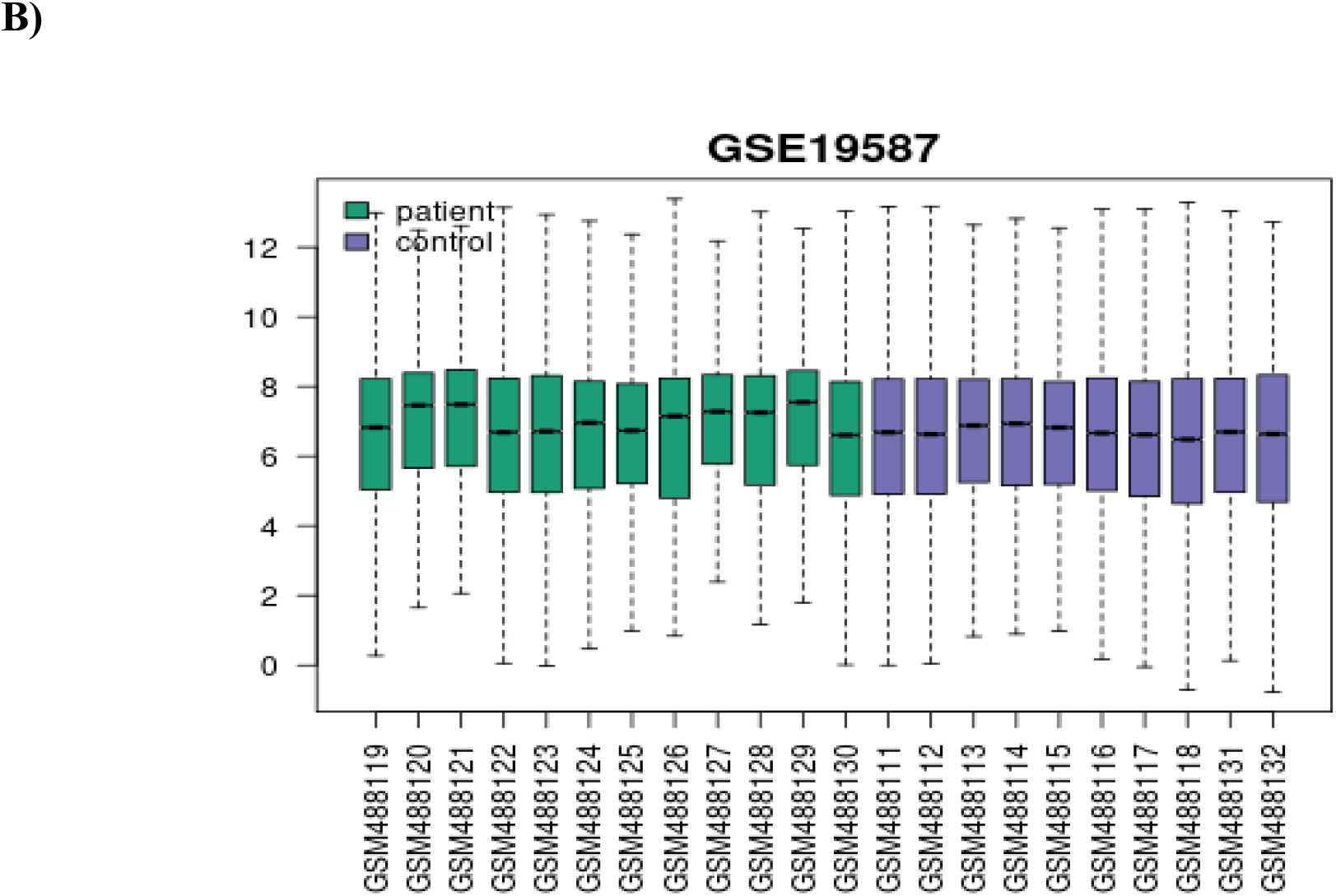
Box plots of A) Glioblastoma (GSE31262) and B) PD (GSE19587) representing the normalization data. After data normalization with the GEO2R program, the distribution of patients versus controls was determined.

**Fig. -2:**
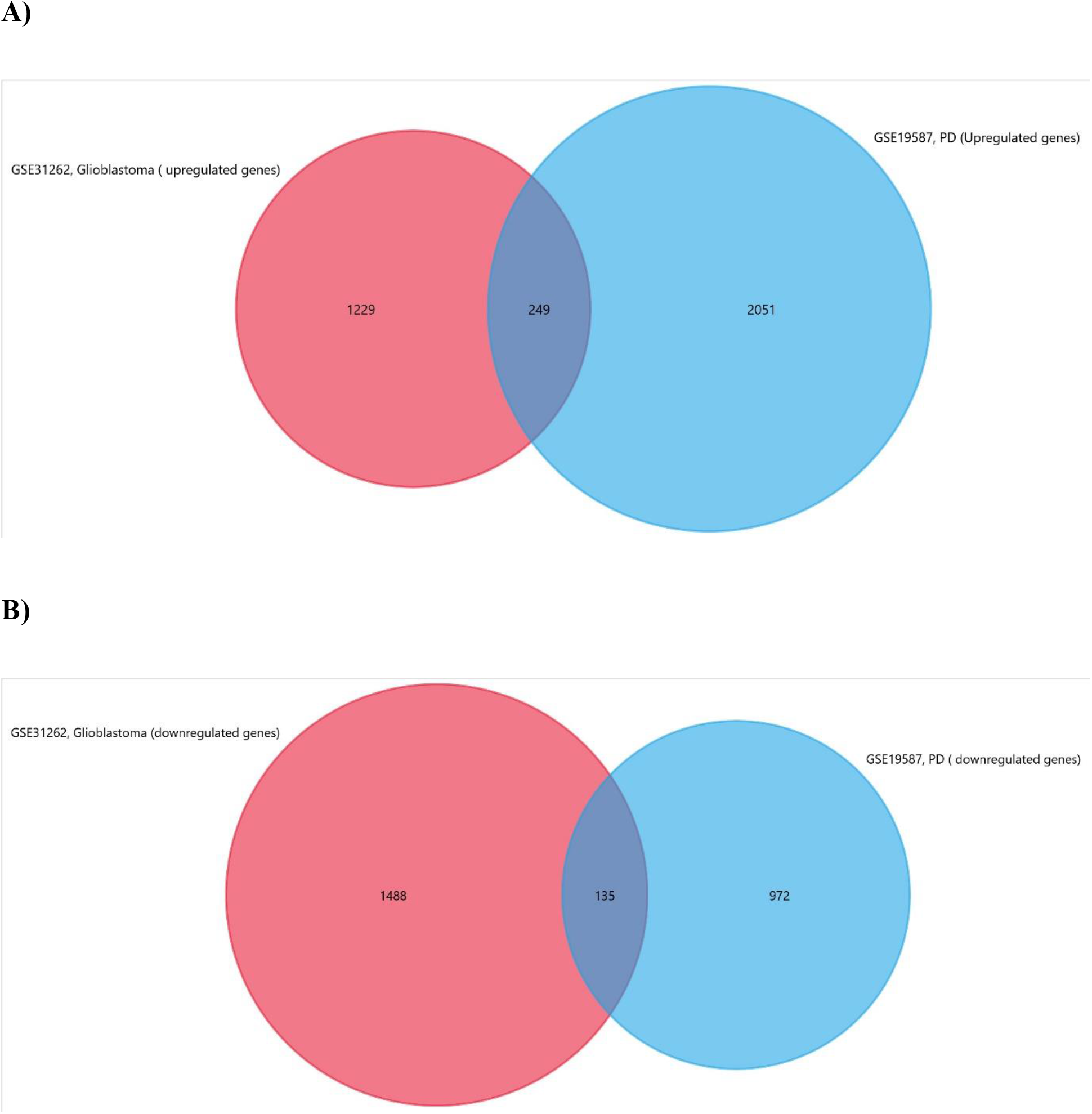
Venn diagram representing the DEGs among the two datasets, GSE31262 for glioblastoma and GSE19587 for PD. A) common upregulated DEGs, B) common downregulated DEGs in these diseases.

After that we imported 384 common DEGs, both upregulated and downregulated into the STRING tool and obtained the protein-protein interaction (PPI) network. We concealed The disconnected genes by clicking ‘hide disconnected nodes in the network’ inside the ‘network analysis’ option and selected the ‘medium confidence option’ in the ‘minimum required interaction score option’ in “Settings”. Total 382 nodes and 1468 edges were present in the protein-protein interaction network. The average node degree is 7.69. Therefore we downloaded the PPI as a high-resolution bitmap in PNG format (Figure-3).

**Fig. -3:**
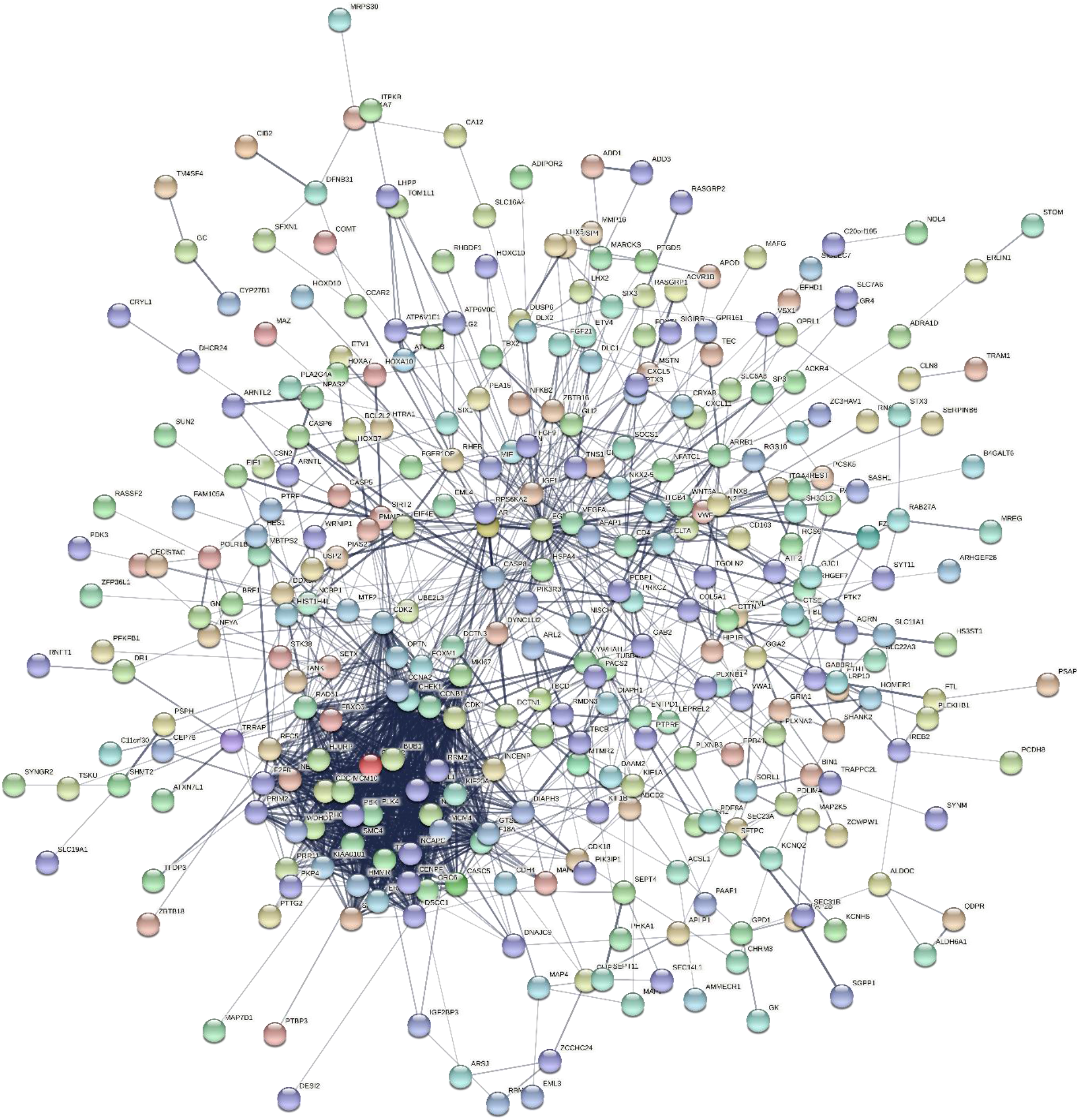
STRING protein-protein interaction network of common DEGs including common upregulated DEGs and common downregulated DEGs. In this network, there are 382 nodes and 1468 edges. Lines represent interaction and the circles represent common genes. The color of the lines represents the type of interaction.

To find out the hub genes, the whole protein-protein interaction network was subjected to Cytoscape directly from the STRING tool. The CytoHubba plugin was used within the Cytoscape software to obtain the top 10 hub genes by setting the confidence (score) cut-off at 0.40 and the maximum number of additional interactions to 0 (Figure-4).

**Fig. -4:**
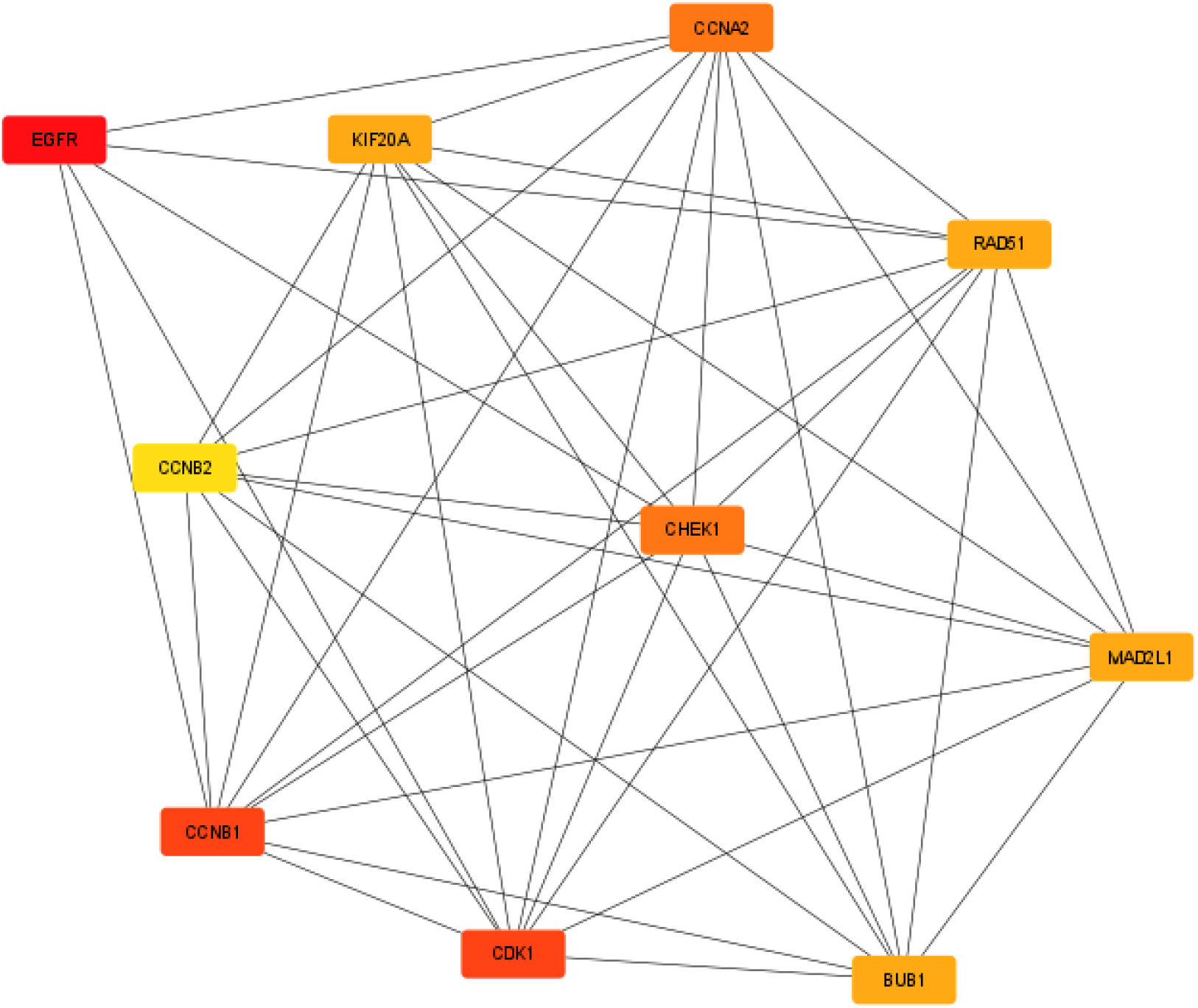
Network of the top ten hub genes Using Cytoscape software. The color represents the degree of connectivity; red represents the highest degree, orange represents the intermediate degree, and yellow represents the lowest degree.

It was found that Epidermal Growth Factor Receptor (EGFR, degree=56) possessed the highest degree that’s mean it shows the highest connectivity with other hub genes and it is followed by Cyclin B1 (CCNB1) and Cyclin-Dependent Kinase 1(CDK1) with a degree of 54, Cyclin A 2 (CCNA2) and Checkpoint Kinase 1 (CHEK1) had a degree of 51, RAD51 Recombinase (RAD51), Mitotic Arrest Deficient 2 Like 1 (MAD2L1), Kinesin Family Member 20A (KIF20A), Bub 1 Mitotic Checkpoint Serine/Threonine Kinase (BUB1) all had a degree of 47, and Cyclin B2 (degree=45) possessed the lowest degree (Table-2). These top 10 hub genes were upregulated in these diseases, no downregulated genes were found among the hub genes.

**Table-1:** Dataset information obtained from GEO. There are 9 patient samples and 5 control samples in dataset GSE31262 and 12 patient samples and 10 control samples in dataset GSE19587.

| Diseases | Datasets | No. of patients | No. of Controls |
| --- | --- | --- | --- |
| **Glioblastoma** | GSE31262 | 9 | 5 |
| **PD** | GSE19587 | 12 | 10 |
| **Total** |  | 21 | 15 |

**Table-2:** Top 10 hub genes and their degree.

| Hub Genes | Degree |
| --- | --- |
| EGFR | 56 |
| CCNB1 | 54 |
| CDK1 | 54 |
| CCNA2 | 51 |
| CHEK1 | 51 |
| RAD51 | 47 |
| MAD2L1 | 47 |
| KIF20A | 47 |
| BUB1 | 47 |
| CCNB2 | 45 |

The top 10 hub genes were then imported to miRNet software to determine their interactions with miRNAs. After that miRNet created a network between common DEGs, miRNAs, and transcription factors. Total 581 miRNAs, 63 transcription factors were found and they made the interactions with 10 genes. In this network total of 1054 edges were found (Figure-5).

**Fig. -5:**
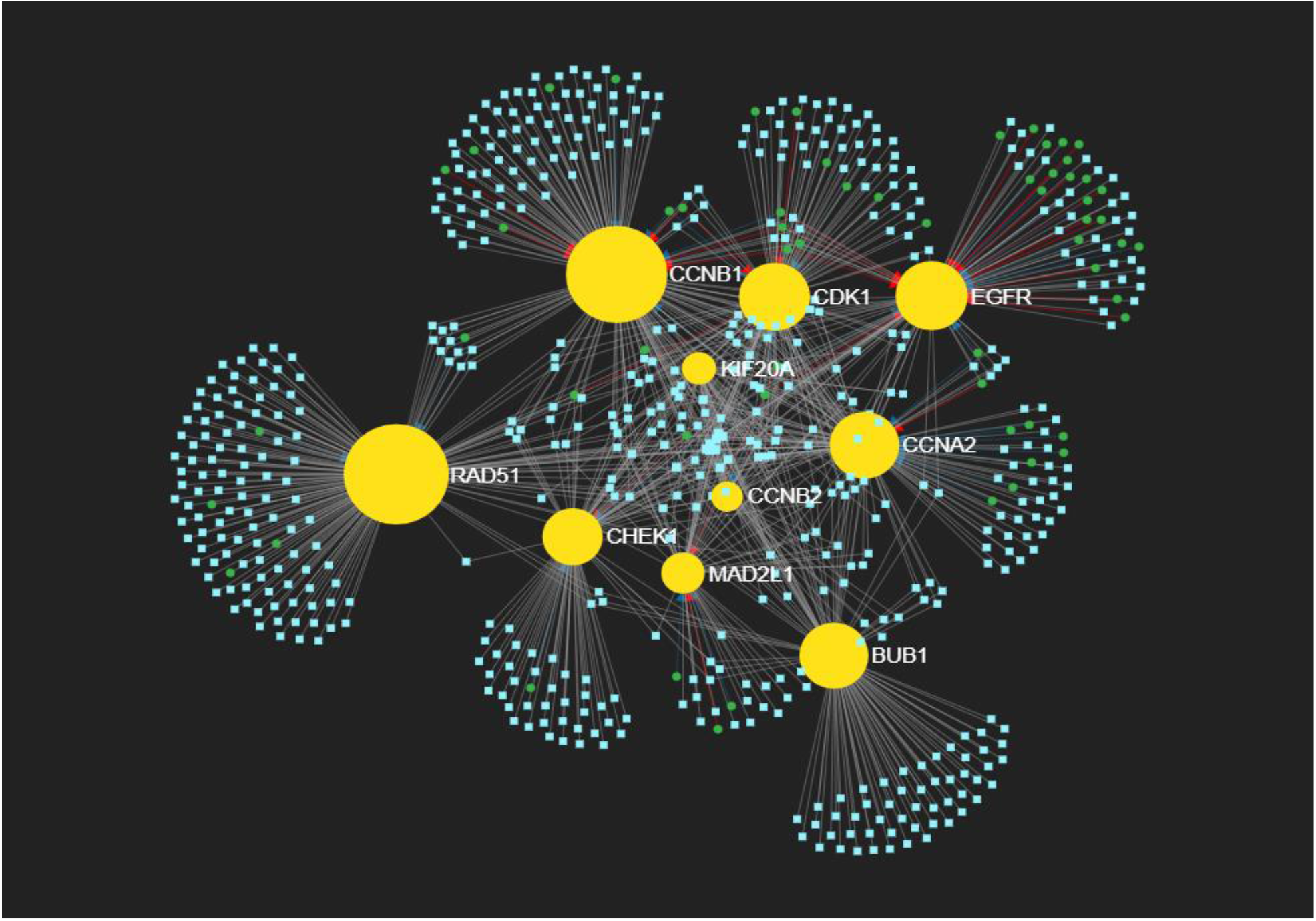
The interactions between the top ten hub genes and the microRNAs, transcription factors are shown. Sky-blue color represents the micro RNAs, yellow represents the top ten hub genes, and green color represents transcription factors.

We found that CCNB1(degree=186) had the highest connectivity and was followed by RAD51 with a degree of 168, CCNA2 with a degree of 127, CDK1 with a degree of 120, EGFR with a degree of 113, BUB1 with a degree of 101, CHEK1 with a degree of 95, MAD2L1 with a degree of 61, KIF20A with a degree of 49 and CCNB2 with a degree of 34. We also identified that there were four miRNAs with the highest connectivity. They were hsa-mir-16-5p, hsa-mir-34a-5p, hsa-mir-124-3p, and hsa-mir-147a with a degree of 10 and followed by hsa-mir-103a-3p, hsa-mir-107, and hsa-mir-129-2-3 had a degree of 9, hsa-mir-23b-3p and hsa-mir-195-5p with a degree of 8 and hsa-mir-205-5p with a degree of 7 (Table-3).

**Table-3:** List of top 10 hub genes and miRNAs with their degree.

| **Name of DEGs and miRNAs** | **Degree** |
| --- | --- |
| CCNB1 | 186 |
| RAD51 | 168 |
| CCNA2 | 127 |
| CDK1 | 120 |
| EGFR | 113 |
| BUB1 | 101 |
| CHEK1 | 95 |
| MAD2L1 | 61 |
| KIF20A | 49 |
| CCNB2 | 34 |
| hsa-mir-16-5p | 10 |
| hsa-mir-34a-5p | 10 |
| hsa-mir-124-3p | 10 |
| hsa-mir-147a | 10 |
| hsa-mir-103a-3p | 9 |
| hsa-mir-107 | 9 |
| hsa-mir-129-2-3 | 9 |
| hsa-mir-23b-3p | 8 |
| hsa-mir-195-5p | 8 |
| hsa-mir-205-5p | 7 |

After that, we imported the common DEGs into an online software, Enrichr to analyze the GO ontology such as GO biological processes (GO-BF), GO molecular functions (GO-MF), GO cellular components (GO-CC), and the KEGG pathways associated with these common DEGs. The enriched biological processes are mitotic nuclear membrane disassembly, nuclear membrane disassembly, regulation of cyclin-dependent protein serine/threonine kinase activity, G2/M transition of mitotic cell cycle, cell cycle G2/M phase transition, negative regulation of mitotic cell cycle, mitotic nuclear membrane organization, mitotic nuclear membrane reassembly, nuclear membrane reassembly and regulation of cyclin-dependent protein kinase activity (Figure-6A). The enriched molecular functions are cyclin-dependent protein serine/threonine kinase regulator activity, protein kinase regulator activity, kinase binding, protein serine/threonine kinase activity, protein kinase binding, patched binding, histone threonine kinase activity, histone kinase activity, cyclin-dependent protein serine/threonine kinase activator activity, and MAP kinase kinase kinase activity (Figure-6B). The enriched cellular components are cyclin-dependent protein kinase holoenzyme complex, serine/threonine-protein kinase complex, intracellular membrane-bounded organelle, nucleus, condensed nuclear chromosome, spindle, condensed chromosome, nuclear chromosome, mitotic spindle, and cyclinA2-CDK2 complex (Figure-6C). The KEGG pathways, which are associated with these common DEGs, are cell cycle, progesterone-mediated oocyte maturation, oocyte meiosis, cellular senescence, p53 signaling pathway, human immunodeficiency virus 1 infection, human T-cell leukemia virus 1 infection, foxo signaling pathway, viral carcinogenesis, and pancreatic cancer (Figure-6D).

**Fig. -6:**
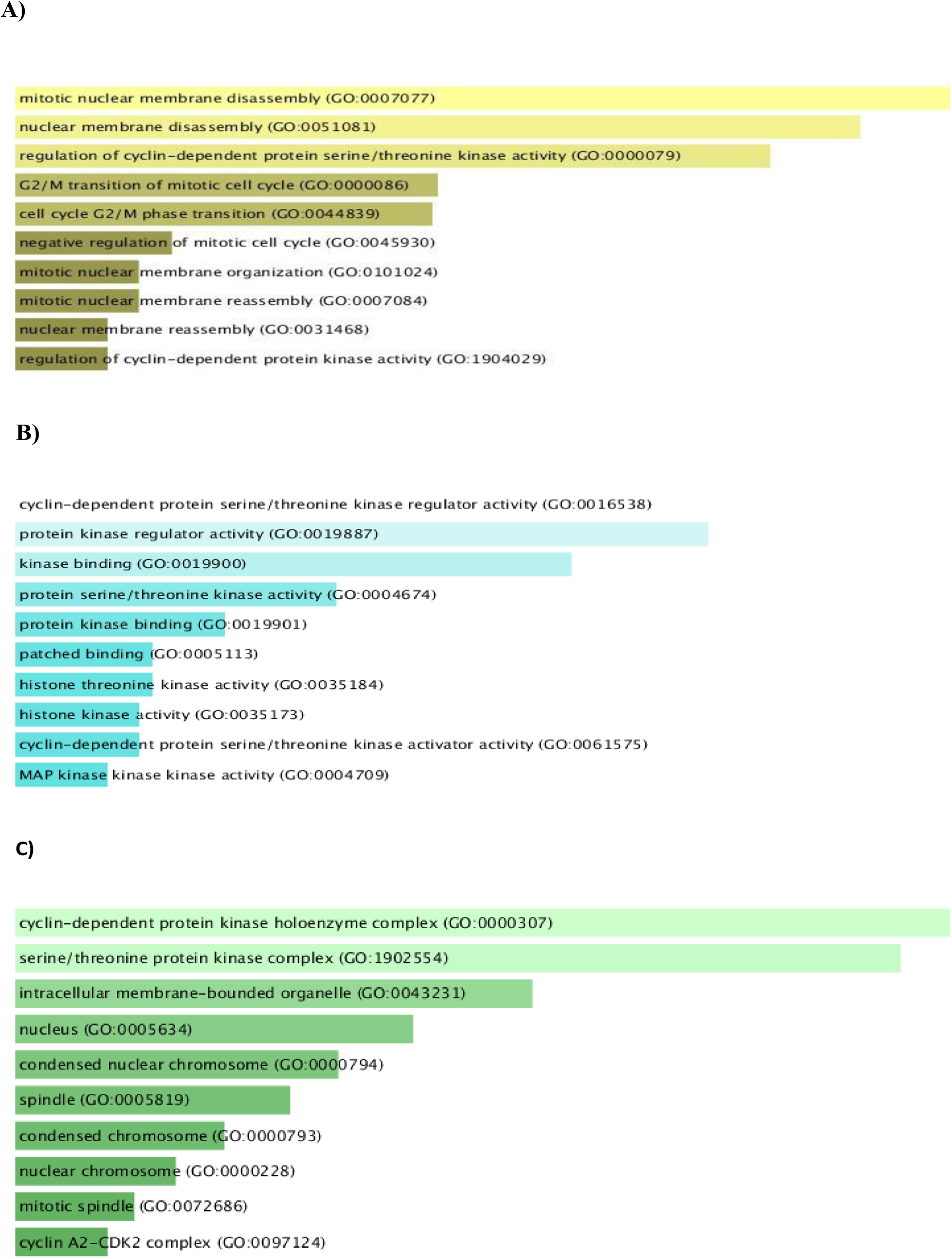
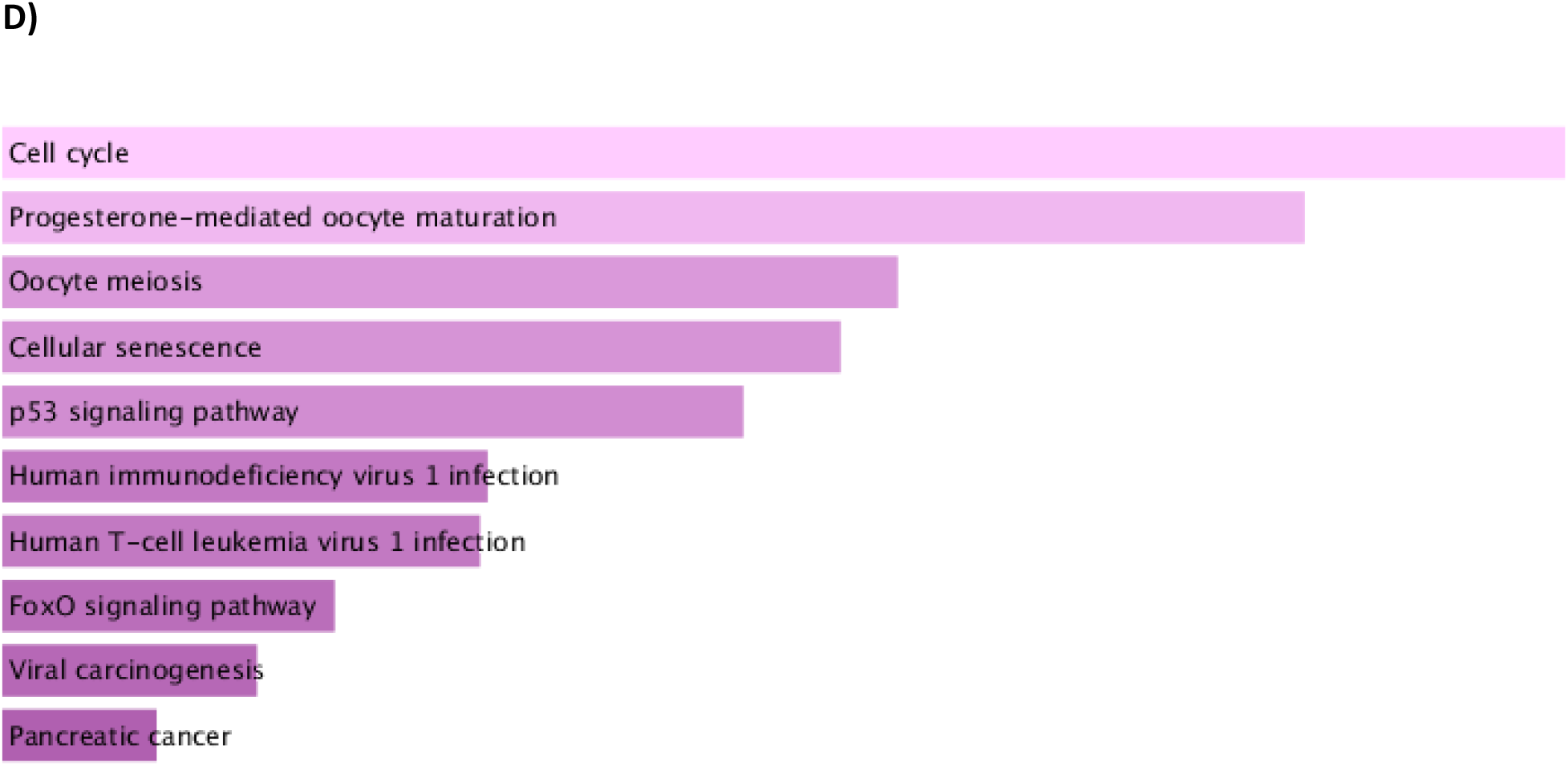
Graphical representation of enrichment analysis. A) Top 10 enriched biological processes in hub genes. The x-axis represents several genes, and the y axis represents the biological processes. B) Top 10 enriched molecular functions in hub genes. The x-axis represents the number of genes, and the y axis represents the molecular functions. C) Top 10 enriched cellular components in hub genes. The x-axis represents the number of genes, and the y axis represents cellular components. D) Top 10 enriched KEGG pathways in hub genes. The x-axis represents the number of genes, and the y axis represents the KEGG pathways.

## 4. Discussion

In this study, we wanted to investigate the differentially expressed genes and their similarities in these diseases as well as the common DEGs in these diseases, miRNAs association with the hub genes and the biological processes, molecular functions, cellular components, and the pathways associated with these two diseases; Glioblastoma and Parkinson’s disease.

The top hub genes, that were found by us in these two datasets, are EGFR, CCNB1, CDK1, CCNA2, CHEK1. In a study, it was seen that the overexpression, amplification, and mutation of the EGFR cause approximately 40 to 50 percent of tumors to develop into Glioblastoma (Hatanpaa et al., 2010). A very interesting observation was made in another study. It was found that in the early stage of Parkinson’s disease (PD), there was a reduction in plasma EGFR level, whereas, in the advanced stage of PD, EGFR plasma level increased (Jiang et al., 2015). Cyclin B1(CCNB1)-CDK1 complex, cyclin B2(CCNB2)-CDK1 complex regulate the G2/M phase of cell cycle so CDK1 abnormality can cause different types of cancer in the human body and it is found that CCNB1, CCNB2, CDK1 are three of the hub genes associated with hepatocellular carcinoma (Zou et al., 2020). CDK1 also plays a key role in glioblastoma development along with CDK2 and TOP2A (Bo et al., 2017). Some hub genes, (discovered in glioblastoma and PD datasets) such as CDK1, CCNB1, CCNA2, MAD2L1, BUB1 are also involved in the progression of lung adenocarcinoma (Chen et al., 2020). In another study, we found the involvement of CHEK1 in glioblastoma but its molecular and biological functions in glioblastoma remain unspecified still now (Bai et al., 2018). But the roles of some hub genes in Parkinson’s disease are still unclear.

We also studied the interactions between the hub genes and miRNAs. From the network, which is obtained by the miRNet software, we found that CCNB1 had the highest connectivity with maximum microRNAs and transcription factors. MicroRNAs such as hsa-mir-16-5p, hsa-mir-34a-5p, hsa-mir-124-3p, hsa-mir-147a had the highest connectivity with a degree of 10. In one study, hsa-mir-195-5p (degree=8) was found to be upregulated in glioblastoma patients (Géczi et al., 2021). But in another study, it was seen that miR-16-5p was downregulated in glioblastoma (Krell et al., 2018). We also found from a study that hsa-mir-16-5p(degree=10) and hsa-mir-23b-3p (degree=8) act as potential biomarkers in PD (Taguchi & Wang, 2018). The overexpression of hsa-mir-124-3p miRNA leads to the inhibition of glioblastoma ‘cell proliferation’, ‘migration’ and finally results in glioblastoma ‘cell arrest’ and ‘cell apoptosis’ (Zhang et al., 2018) while some studies showed it to be downregulated in ALS patients and one study showed it to be upregulated (Foggin et al., 2019).

## 5. Conclusion

Although each disease has been studied separately in the past, these two diseases (glioblastoma and Parkinson’s disease) have never been studied together before this. We created a PPI network of common up and downregulated DEGs and identified the top ten hub genes and their relationships with microRNAs. There is a chance that these hub genes can be used as drug targets. More research is needed to validate the above findings.

## Supporting information

GEO accession number

Hub genes with their degree

hub genes and microRNA with degree

